# Bioassemblying Macro-Scale, Lumnized Airway Tubes of Defined Shape via Multi-Organoid Patterning and Fusion

**DOI:** 10.1101/2020.11.01.363705

**Authors:** Ye Liu, Catherine Dabrowska, Antranik Mavousian, Bernhard Strauss, Fanlong Meng, Corrado Mazzaglia, Karim Ouaras, Callum Macintosh, Eugene Terentjev, Joo-Hyeon Lee, Yan Yan Shery Huang

## Abstract

Epithelial, stem-cell derived organoids are ideal building blocks for tissue engineering, however, scalable and shape-controlled bioassembly of epithelial organoids into larger and anatomical structures has yet to be achieved. Here, a robust organoid engineering approach, Multi-Organoid Patterning and Fusion (MOrPF), is presented to assemble individual airway organoids of different sizes into upscaled, scaffold-free airway tubes with pre-defined shapes. Multi-Organoid Aggregates (MOAs) undergo accelerated fusion in a matrix-depleted, free-floating environment, possess a continuous lumen and maintain prescribed shapes without an exogenous scaffold interface. MOAs in the floating culture exhibit a well-defined three-stage process of inter-organoid surface integration, luminal material clearance and lumina connection. The observed shape stability of patterned MOAs is confirmed by theoretical modelling based on organoid morphology and the physical forces involved in organoid fusion. Immunofluorescent characterization shows that fused MOA tubes possess an unstratified epithelium consisting mainly of tracheal basal stem cells. By generating large, shape-controllable organ tubes, MOrPF enables upscaled organoid engineering towards integrated organoid-devices and structurally complex organ tubes.

## 1. Introduction

Engineered fusion of multi-cellular materials, such as spheroids and organoids, represents a biomimetic cell assembly process fundamental to the fields of biofabrication, tissue engineering and *in vitro* tissue modelling^[1],[2],[3]^. Organization and fusion of cell aggregates to achieve prescribed shapes (and functions) depends on the selection of initial multi-cellular building blocks. Coalescence of solid-core spheroids can readily occur via so-called ‘tissue liquidity’^[4]^, in small aggregates of epithelial cells^[1]^, fibroblasts^[5]^, mesenchymal stem cells^[6]^, and in hybrid spheroids consisting multiple cell types^[7],[8]^. Guided-assembly through micro-fabricated molds or 3D-printing can further define the three-dimensional (3D) architecture of engineered tissues^[9]^. However, due to the diffusion limit, fused tissue constructs from solid-core spheroids are hardly upscaled to viable thick tissues beyond millimeter scale, without additional perfusion platforms or engineered vasculatures^[3]^. In the last decade, a class of self-organizing multi-cellular material called organoids, have emerged as a powerful tool to study the behavior of their tissue of origin^[10],[11],[12]^. Given their close resemblance to native organs in histology and cell composition, organoids represent ideal modular units for the biofabrication of biomimetic organs^[13],[14]^. While fusion has been demonstrated in pairs of brain organoids^[15],[16]^ and in collagen-embedded intestinal organoids^[17]^, the resulted fused products do not exhibit organ-scaled size and anatomical shape. Upscaled fusion of cystic epithelial organoids into shape-controllable, large lumenized tissues is challenging. This is because to produce large, functional epithelial tubes with an elongated lumen, such as the trachea, fusion of organoid cysts requires not only the seamless surface integration between adjacent organoids but also the inter-connection of the fluid-filled lumina.

Here, using mouse tracheal epithelial organoids, we show multi-organoid patterning and fusion (MOrPF), to create lumenized, macro-scale organ tubes of defined shapes. We patterned individual organoids into shape-defined multi-organoid aggregates (MOAs) and employed a free-floating environment to encourage inter-organoid surface connection and lumenization. Notably, the size of fused organ tubes can be prescribed to match that of an adult mouse trachea. A theoretical model is proposed to explain MOA shape maintenance post-patterning, in the absence of a solid matrix or shape-supporting scaffold interface. We envisage these MOA tubes as a foundation for several emerging downstream applications, including organoid-microfluidics integration and multi-tissue organ reconstruction.

## 2. Results

### 2.1 MOAs undergo a three-stage fusion process leading to lumenized tube formation

Starting with individual airway organoids derived from adult mouse tracheal epithelial cells, we established a robust organoid fusion platform, for the upscaled engineering of size-relevant epithelial organ tubes (**Figure 1**a). Isolated single tracheal basal stem cells form cystic airway organoids of heterogeneous sizes in 3D Matrigel (a mouse-derived extracellular matrix), as previously reported^[18]^ (**Figure S1**a, Supporting information). Occasionally, we observed pairs of organoids fused into randomly shaped cysts (Figure S1b, Supporting information). To test whether liberating organoids from Matrigel would enhance fusion, we mechanically dissociated airway organoids from Matrigel, and cultured them in a free-floating condition (termed ‘Matrigel-depleted culture’). Though a low amount of Matrigel could be carried over into the MOrPF process, this should not be enough to form a constraining layer around organoid surfaces. In such Matrigel-depleted condition, organoids fused more frequently and formed inter-connected lumina (Figure S1c-d, Supporting information); however, the size and shape of fused products remained heterogeneous. To control the final geometry of fusion products and to improve overall fusion efficiency, we developed a multi-organoid patterning and fusion (MOrPF) workflow as shown in Figure.1a-II. Day-12 organoids were dissociated from Matrigel drops and transferred in a 3D-designed polydimethylsiloxane (PDMS) mold to form shape-patterned multi-organoid aggregates (MOAs). Depending on the PDMS template size and airway organoid diameter, approximately 100-1400 organoids were assembled in each well of the PDMS mold. Subsequently, MOAs were released from the PDMS template and cultured in the floating condition for effective fusion and lumenization.

**Figure 1.**
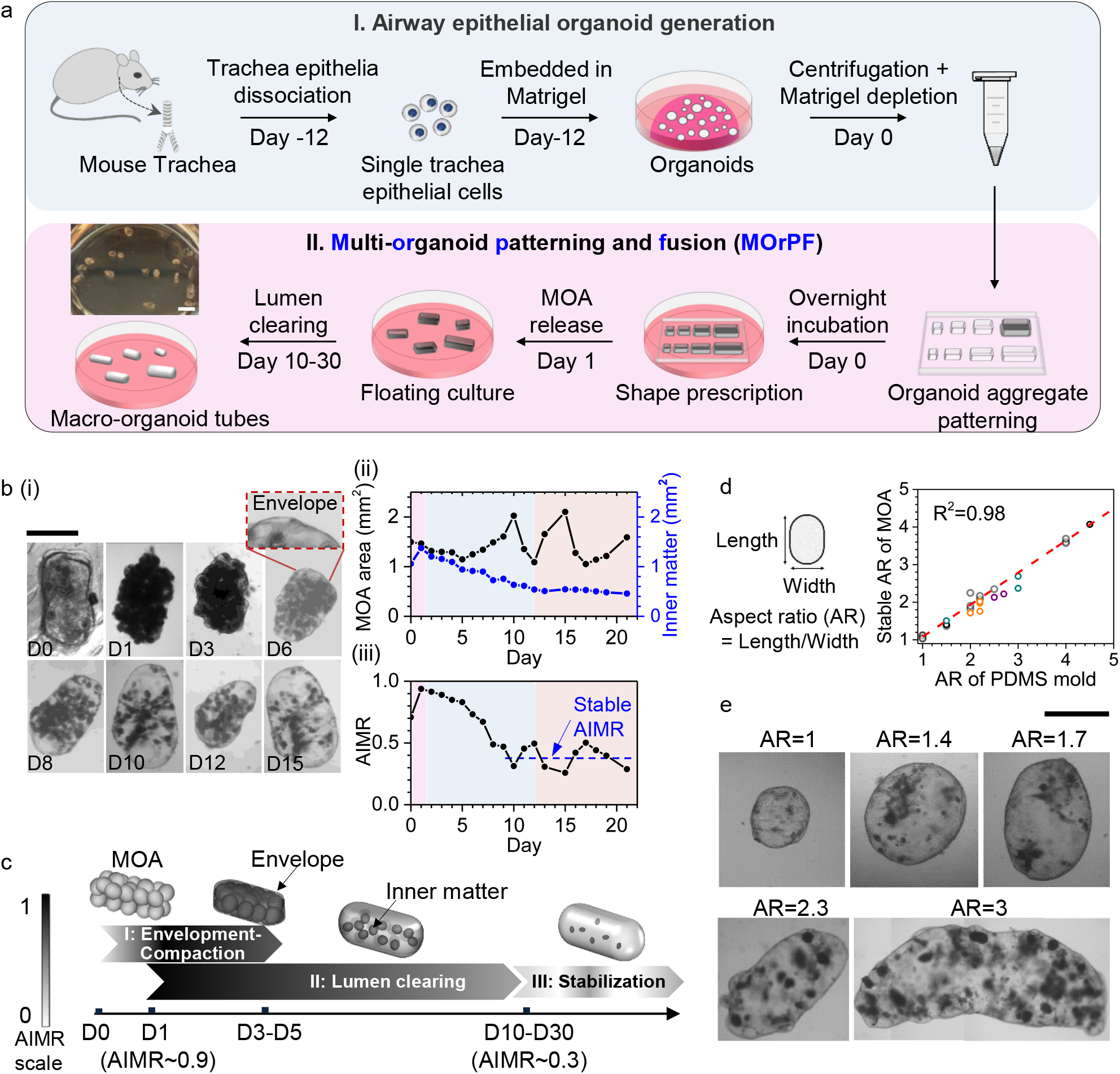
MOrPF technique guides the spatial assembly and fusion of airway organoids into upscaled, lumenized epithelial tubes. a, Schematic representation of mouse airway epithelial organoid generation (I), and the MOrPF procedure (II). Scale bar, 4 mm. b, (i) Representative image sequences (4 independent experiments) of a developing MOA. The MOA was patterned in a PDMS well on Day 0 and released into the floating culture on Day 1. Scale bar, 1mm; (ii) Dynamics of MOA fusion represented by the projected MOA area (black curve) and the inner matter area (blue curve); (iii) quantification of AIMR dynamics, approximating into a three-stage process. AIMR=MOA area/Inner matter area. c, Schematic representation of the three stages in MOA fusion. MOA grey value corresponds to the AIMR scale. d, Aspect ratio of stabilized MOAs as a function of their initial aspect ratio prescribed by the PDMS mold. n=26 samples, from 6 independent experiments (each represented with a different color symbol). e, Representative images (4 independent experiments) of engineered MOA tubes with a range of sizes and shapes. Scale bar, 1 mm.

We quantified the overall morphological changes of MOAs during the MOrPF process, by measuring the projected area of the MOA in relation to its inner dense matter (cellular materials in the center of organoids and MOAs), across different stages of MOrPF (Figure 1b, **Figure S2**a, Supporting information). Upon release from the PDMS mold, the patterned MOA retained its prescribed geometry with an opaque appearance (Day 1). An external layer of cells started to envelop the outer surface of the MOA (Day 3, envelopment), leading to the smoothening of MOA external contour by day 6. During this time, the luminal content of the MOA gradually cleared out (Figure 1b(ii)), as quantified by the MOA opacity parameter, i.e. the Apparent Inner Matter Ratio (AIMR) (Figure 1b(iii)). By Day-10, the MOA resembled a quasi-translucent, closed-end tube, which exhibited intermittent lumen shrinkage and re-expansion (Figure 1b(ii)). Similar epithelium rupture and inner fluid release, followed by lumen closure and re-growth, have been observed in single epithelial organoid^[19]^ and the mouse blastocytes^[20]^.

Since the time required for MOA fusion increases with the increase in MOA size (Figure S2f, Supporting information), we chose AIMR, instead of days in culture, to normalize the fusion stages among MOAs of different sizes. By correlating AIMR dynamics with bright field images, we identified three critical stages during MOA fusion progression: Stage I: Envelopment and Compaction; Stage II: Lumen Clearance; and Stage III: Stabilization (Figure 1c). These key stages were typical for patterned MOAs of a range of sizes (Figure S2b-d, Supporting information). During Stage I, MOAs acquired an outer layer of enveloping cells and underwent compaction, showing a peak AIMR value ~0.9 (Figure 1b(iii), Figure S2d, Supporting information). During Stage II (the longest stage in MOrPF), MOAs gradually cleared their luminal content, resulting in a decrease in AIMR from ~0.9 to ~0.35 (Figure 1b(iii), Figure S2d-e, Supporting information). From Stage III, stabilized MOAs developed a smooth outline and a quasi-translucent lumen, while retaining the aspect ratio prescribed by the PDMS mold (Figure 1d-e). Together, our data suggest that MOrPF is a robust process for the assembly and fusion of small, heterogeneously-sized organoids into macro-scale, shape-definable epithelial tubes.

### 2.2 Inter-organoid envelopment during Stage I is a pre-requisite for MOA fusion

We next investigated organoid behaviors throughout the MOA fusion process, by live imaging small clusters of organoids cultured under different conditions. We first observed organoid aggregation and cellular bridge formation (termed ‘inter-organoid envelopment’) between adjacent, touching organoids in the floating culture (**Figure 2**a(i)). After dissociation from Matrigel, those organoids were able to aggregate spontaneously. Upon establishment of the inter-organoid contact, a cellular bridge started to form between the touching organoids and continued to expand into a shared envelope covering the organoids (Figure 2a(ii), Video S1, Supporting information). Within a few days, inter-organoid envelopes became visible at multiple locations of a MOA, gradually integrating into a continuous layer encasing all organoids within the aggregate (**Figure S3**a, Supporting information). Interestingly, organoid aggregation and envelopment was significantly reduced with increased Matrigel concentration, as supported by our comparative data in Gel-suspension (Matrigel added at a 20% volume concentration in medium, Figure 2b, Figure S3b, Supporting information) and 100% Matrigel-embedded cultures (Figure 2c, Figure S3d-e, Supporting information). In the Gel-suspension condition, inter-organoid enveloping efficiency dropped to below 30% (Figure 2e) and the onset of envelopment was delayed after organoids had established contact (Figure 2b(ii)). When embedded in 100% Matrigel, organoids rarely aggregated or developed envelopes, even in close proximity (Figure 2c-e). Without efficient envelopment, MOAs failed to develop fused lumina (Figure 2f). Fusion efficiency in MOAs (with projected area of 0.6-1 mm^2^) was <20% in Gel-suspension and <10% in Matrigel-embedded culture (Figure 2f). By contrast, in the floating condition, over 60% of organoids formed lumenized MOAs by Day 7, which further increased to over 90% by Day 11. To further explore organoid fusion capability in the presence of a nonadherent hydrogel such as agarose, we placed organoids in a floating culture supplemented with 0.5% weight percent agarose. Organoid aggregation and envelopment were not affected by agarose fragments, but the lumen clearing process seemed restricted (Figure S3f, Supporting information). Together, these data suggest that envelope formation is a pre-requisite for organoid fusion and lumen connection. Inter-organoid envelopment is enhanced in the floating culture devoid of adhesive Matrigel matrix, which largely liberates organoids from organoid-substrate interaction, while permitting organoid-organoid interaction.

**Figure 2.**
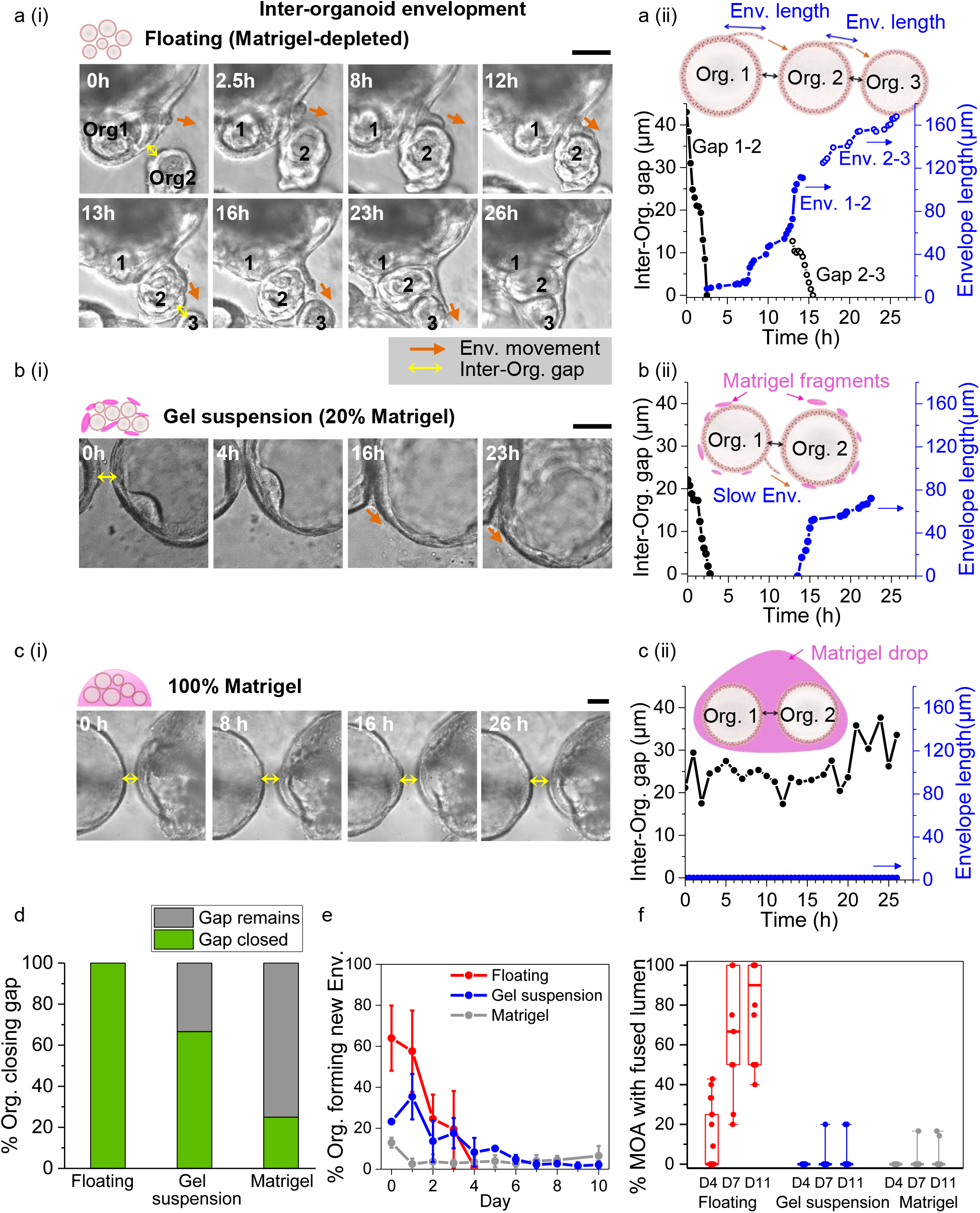
Inter-organoid envelopment is a pre-requisite for MOA fusion. a-c, Live imaging and quantification of inter-organoid envelope dynamics in Floating (a), Gel suspension (b) and full Matrigel (c) cultures. a(i), b(i), c(i), Representative image sequences (3 independent experiments) highlighting changes in inter-organoid gaps and the envelope leading edge positions for the three culture systems. Scale bars, 100 μm. a(ii), b(ii), c(ii), Measurement of inter-organoid gaps and envelope lengths for the corresponding organoids shown in the left panel. d, Percentage of organoids closing inter-organoid gaps (less than 80 μm) in the three culture systems. Floating, n=20; Gel suspension, n=15; Matrigel, n=20. e, Percentage of closely-spaced organoids developing new envelopes daily in the three cultures. Floating, n=193; Gel suspension, n=331; Matrigel, n=449. Data are presented as mean ± SD. f, Percentage of closely-spaced organoids undergoing successful fusion (as marked by the formation of a continuous lumen) in different culture systems. Midline = median, box = 25th-75th percentiles, Whisker = min and max values. Floating, n=10; Gel suspension, n=17; Matrigel, n=17.

### 2.3. Cellular matter release results in MOA lumen clearance

After the establishment of a shared envelope, we further observed MOA compaction and inner matter release, using live imaging on MOAs with a 2-day interval, over 8 days. Coinciding with envelope extension, inter-organoid space decreased within MOAs (**Figure 3**a), allowing increased interfacial contact between adjacent organoids (Video S2, Supporting information). During the lumen clearing stage, MOAs developed large, connected cavities by releasing their inner contents (termed ‘inner matter release’) from multiple epithelium rupture sites (Figure 3b(i), Video S3-4, Supporting information). Given that the AIMR of MOAs gradually stabilized by the end of Stage II-lumen clearance, we asked whether the local release of inner matter correlated with the ensemble-level MOA opacity. Following the local release dynamics of MOAs in their early (AIMR>0.8), mid (0.4<AIMR<0.7) and late (AIMR<0.4) lumenization phases, we found that MOAs extruded inner matter in an intermittent, ‘start and stop’ fashion (**Figure S4**, Supporting information), which could take from a few hours to several days at one release site. We then calculated the local release speed, by quantifying the hourly increase in the projected area of a released inner cluster. As shown in Figure 3b (ii), going from the early, to mid and late lumen clearing phases, both the mean and the maximum release speeds decreased, indicating the accomplishment of lumenization and the stabilization of MOAs.

**Figure 3.**
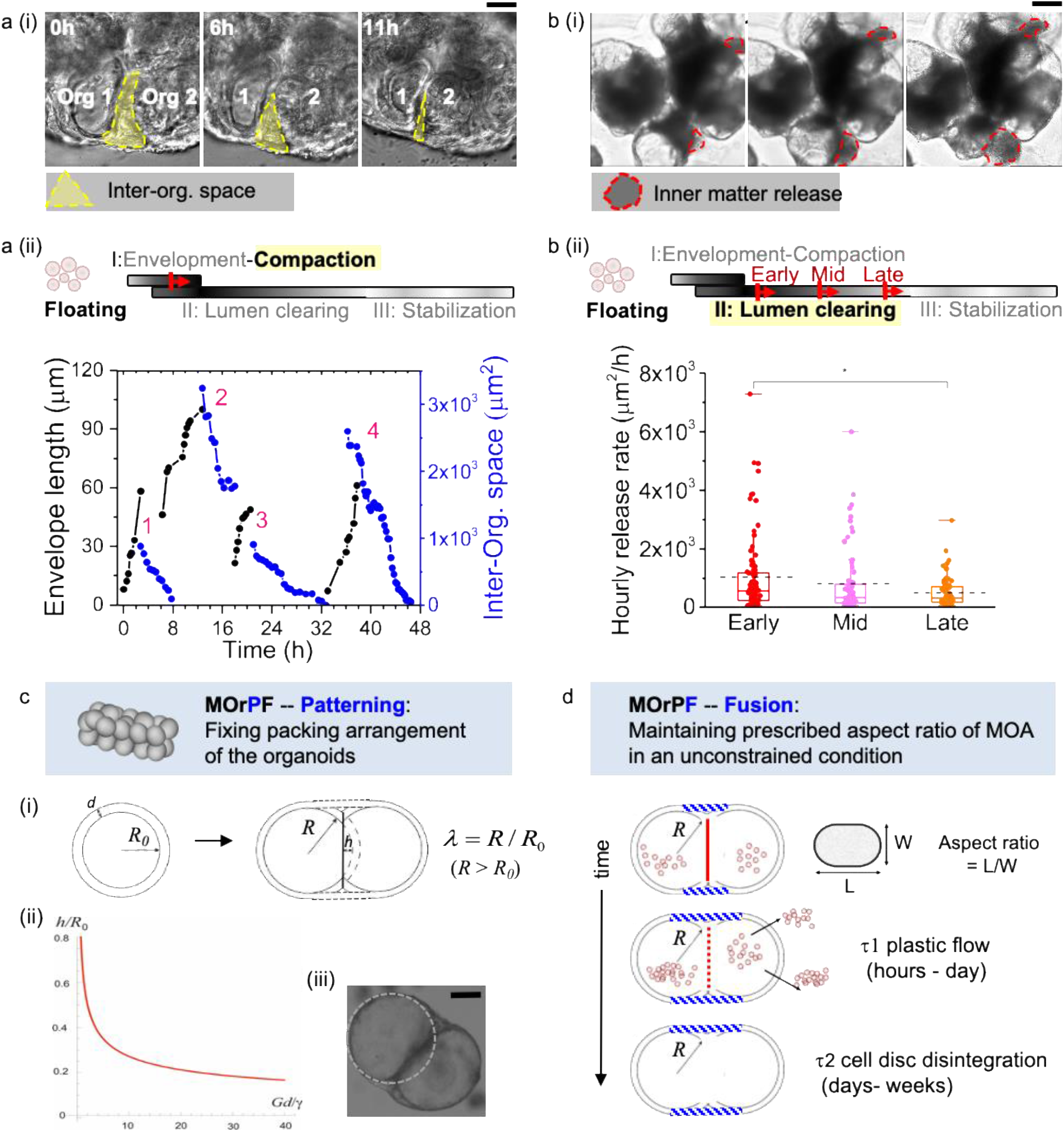
Dynamics of MOA compaction (a) and inner matter release (b) and theoretical accounts for MOA shape stability (c,d) in MOrPF. a(i), Representative image sequence (4 independent experiments) of MOA compaction via the closure of inter-organoid space. Scale bar, 100 μm. a(ii), Measurement of envelope length and the projected area of inter-organoid space over time. n=4 independent samples. Note that upon establishment, envelopes quickly integrated onto the organoid surfaces, with leading edges untrackable in the bright-field images. b(i), Representative image sequence (3 independent experiments) of MOA inner matter release. Scale bar, 500 μm. b(ii), Quantification of the released inner matter per hour in the Early, Mid and Late stages of MOA lumenization. Early, n=79; Mid, n=68; Late, n=70. *P <0.05 by Mann-Whitney U test. c, Schematic representation of a physical model on MOA shape definition in the early stage of MOrPF. (i), Two organoids (radius *R_0_*, shell thickness *d*) fuse to form a dumbbell-like structure consisting of two spherical caps with a larger radius *R*> *R_0_*. *h* denotes how much the two organoids overlap to form the dumbbell. (ii) Theoretical model predicts the overlapping distance *h*/ *R_0_* as a function of *Gd/γ*. (iii) Representative image (2 independent experiments) of two overlapping organoids joint by envelopes. Scale bar, 200 μm. d, Schematic representation of MOA shape maintenance in MOrPF. The dumbbell outer layer fluidizes in τ_1_ (hours to days). After the disintegration of the adhesion disk in τ_2_ (days to weeks), the new dumbbell shape will become the new quasi-equilibrium state, maintained by the high bending rigidity of the epithelial outer layer.

### 2.4. MOA shape stability during fusion can be explained by a theoretical model based on airway organoid morphology and physical forces

Since MOAs retain their prescribed, elongated shapes in the floating culture for weeks (Figure 1d-e), we postulated that such shape stability could originate from the characteristic airway organoid morphology, i.e. fluid-filled cysts enclosed by an epithelial layer (which was treated as an epithelial ‘shell’ in the context of the following physical model). We examined the physical forces at play during the organoid fusion process, the balance of which determines the overall shape of a fused MOA. To construct a simple mathematical model of organoid fusion and MOA shape maintenance, we consider a pair of spherical, equal-sized epithelial organoids (Figure 3c(i)), each of overall thickness *d* (the pseudostratified layer of epithelial cells), filled with incompressible fluid. Before fusion, the organoids (which had grown for 12 days post-plating in Matrigel) are assumed to have a radius *R*_0_. Observations showed that upon their contact, a stable dumbbell-like structure would form (Figure 3c(iii)), consisting of two spherical caps of a new radius *R*, and an interfacial adhesion ‘disk’ in between, which is flat when the two organoids were of the same radius initially. To preserve their inner volume, each organoid shell has to be stretched, which causes an elastic energy penalty^[21],[22]^ in their outer surfaces (see Additional information and **Figure S5**, Supporting information for details). This energy cost, defined by the elastic modulus *G* of the epithelial shell, balances against the energy gain from the adhesion disk due to the favorable surface energy *γ*^23],[24]^. These two competing factors determine the shape of the fused object, before the gradual disintegration of the interfacial disk, which takes a few days to weeks. The preferred shape of the fused organoids is given by the plot of the spherical cap height *h*, as a function of a single control parameter: the non-dimensional ratio *Gd/γ* (Figure 3c(ii)).

Taking the measured values of *G* = 200 Pa ^[21]^ and *d* = 20 μm (**Figure 4**a), and the estimated adhesion energy at the initial cell-cell contact: *γ* = 2 mN/m, we find the control parameter *Gd/γ* = 2, and the predicted height: *h*= 0.5*R*_0_. The observed shape of the organoid dumbbell (Figure 3c(iii)) is almost exactly matching this prediction of *h* ~ 0.5*R*_0_, suggesting that our presented model based on two competing physical forces is a good description of the initial scenario in organoid fusion.

**Figure 4.**
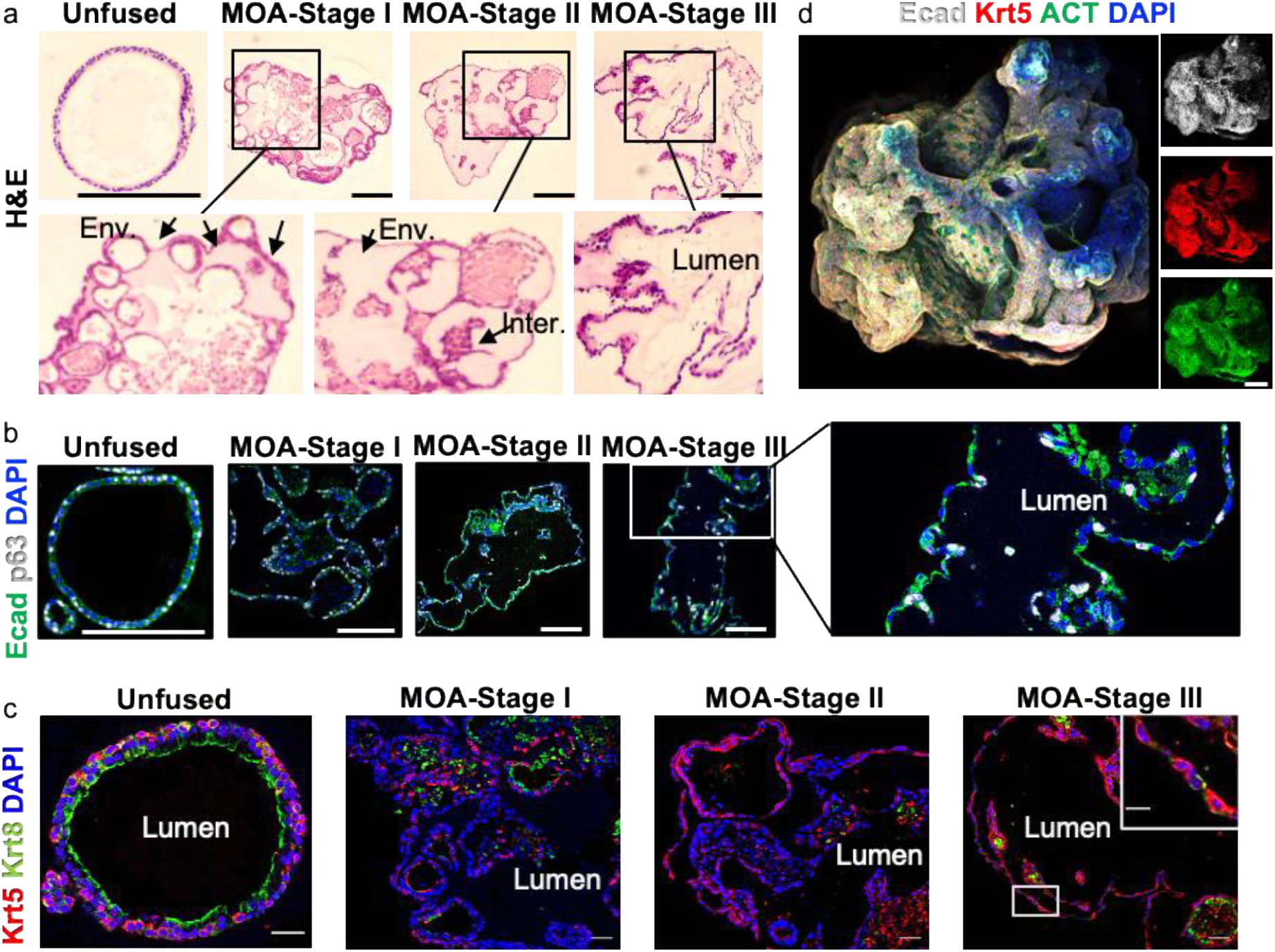
Biological characterization (a-c) and application (d) of bioengineered MOA tubes. a, Representative H&E staining (three independent experiments) of an unfused airway organoid (control) and Stage I to III MOAs. Short arrows indicate ‘envelope’ structures whereas long arrows indicate the remaining ‘interfacial’ disks. Scale bar: 150 μm. b, Representative immunostaining (three independent experiments) of unfused airway organoids and Stage I to III MOAs for epithelial marker Ecad (green) and basal-cell marker p63 (white). Scale bar: 150 μm. c, Representative immunostaining images (two independent experiments) characterize the Stage I-III MOAs for a pseudostratified epithelium compared to unfused control airway organoids. The outer layer of Stage I-III MOAs is a single epithelia of Krt5-expressing cells, intermixed with rare Krt8-expressing cells. Krt5 (red), Krt8 (green) and DAPI (blue). Scale bar, 30 μm. d, 3D projection of a Stage II MOA, stained for nuclei (blue), Ecad (white), Krt5 (red) and ACT (green). Scale bar: 370 μm.

Next, we examined how the dumbbell shape established after the initial inter-organoid envelopment can remain stable over the stages of inner matter release and lumen interconnection. Organoid shell rupture can result in fluid release, and consequently, the release of the weakly bound inner matter material. Such release events equilibrate the external and internal fluid pressure, when the inner matter release stops. Another plausible pressure equilibration event could arise from the cellular plastic flow adjusting the area of the dumbbell shell. With this auto-pressure regulation, the inner fluid pressure would not accumulate excessively even after the disintegration of the interfacial disk. This physical insight could explain the intermittent nature of inner matter release patterns (Figure S4, Supporting information). After the period of plastic flow (hours to days) of cells on the outer shell of fused organoids, the elastic tension will be released, and the internal pressure remains equilibrated. The new dumbbell shape will become the new quasi-equilibrium state, maintained by the high bending rigidity of the shell (see the estimate in Supporting Information) even after the disintegration of the adhesion disk (days to weeks, Figure 3d). Such elongated shape was thus maintained throughout MOrPF, after MOAs being prescribed by the initial patterning process.

### 2.5. Biological characterization of fused MOA tubes

We next assessed the cellular morphology and epithelial composition of MOAs at different stages of the MOrPF process. At Stage I-envelopment and compaction, Hematoxylin and Eosin (H&E) staining revealed that the development of the MOA enveloping cell layer was contributed from the outer epithelial layers of individual organoids positioned near the MOA surface (Figure 4a). Proceeding to Stage II-lumen clearance, individual organoids were mostly unidentifiable, and the contour of MOAs became smooth and continuous. In the MOA center, lumina appeared interconnected though some interfacial discs remained. Finally, in Stage III-stabilization, a continuous lumen developed in MOAs. Immuno-histochemistry staining for airway epithelial markers showed long-term maintenance of airway basal stem cells expressing p63 and Keratin-5 (Krt5) in MOAs up until, and including, Stage III-stabilization (Figure 4b-c). Differentiated ciliated cells expressing acetylated tubulin (ACT) were observed in Stage II MOAs (Figure 4d). However, we observed the gradual loss of differentiated luminal cells expressing Keratin-8 (Krt8), and an increased disorganization of the typical pseudostratified airway epithelium over the fusion process (Figure 4c). To better understand the cellular processes responsible for inter-organoid interactions, we examined key polarity markers during fusion. Whereas unfused organoids displayed intact apical-basal polarization, with the basal membrane expressing fibronectin, the apical junction expressing ZO-1, and the lateral membrane expressing E-cadherin, MOAs revealed disorganized polarization (**Figure S6**). Epithelial cells of stabilized MOAs had reversed polarization, with fibronectin marking the cellular surfaces facing the central lumen. Furthermore, although epithelial cells in the MOAs maintained expression of E-cadherin, expression of ZO-1 was significantly decreased. Our data collectively suggest that upon Matrigel depletion and organoid fusion in the floating condition, fused MOA tubes retain a single epithelial layer of basal cell identity without pseudostratified epithelial morphology, and showed marked changes in epithelial polarity. This is possibly due to the reduction of extra-cellular matrix (ECM) proteins after Matrigel depletion at the outside of MOAs^[25]^.

### 2.6. Downstream processing of MOAs for tissue engineering applications

Since the MOrPF process creates fused airway MOA tubes in a highly efficient and shape-controllable manner, it opens up possibilities for scalable organoid-device integration, multi-tissue interaction, and organ architecture reconstruction. For example, fluid transport is characteristic of many organs and plays an important role in epithelium development and function^[26],[27]^. Our engineered airway MOA tubes, of a size similar to the mouse trachea (1-1.5 mm inner diameter^[28]^), were perfusable and exhibited cyclic lumen expansion and relaxation, in synchronization with a peristaltic input flow of culture medium (**Figure 5**a, Video S5, Supporting information). Such a flow-able system (**Figure S7**a-b, Supporting information) is an initial step for recreating fluid transportation in large engineered organ tubes, where the lumen is accessible within a closed, controllable system separated from the environment on the basal side. It also lays foundation for future studies in epithelial mechanics, as well as for lung-related drug testing and disease modelling^[29]^. Additionally, given that most organ epithelia including trachea epithelium are supported by mesenchymal tissues such as smooth muscle cells (SMCs) and basement membrane, we further engineered a SMC support for fused airway MOA tubes, by co-culturing them with mouse vascular aortic smooth muscle cells (MOVAS). A substrate-detachable SMC sheet was fabricated using a temperature-responsive plate, and wrapped around a fused airway MOA tube to create an integrated tissue (Figure 5b). This proof-of-principle co-culture example illustrates the flexible adaptability of our engineered epithelial tubes for the biofabrication of complex, multitissue organ analogues. Finally, we demonstrated that MOA modular units can be connected to each other and jointly fused into bifurcating organ prototypes, mimicking the hierarchical branching architecture in many epithelial tubes (Figure 5c). The feasibility of fusing mouse intestinal organoids into 4 mm-long intestinal tubes in our floating system further indicated the broader application of MOrPF across other types of epithelial organoids (Figure S7c, Supporting information).

**Figure 5.**
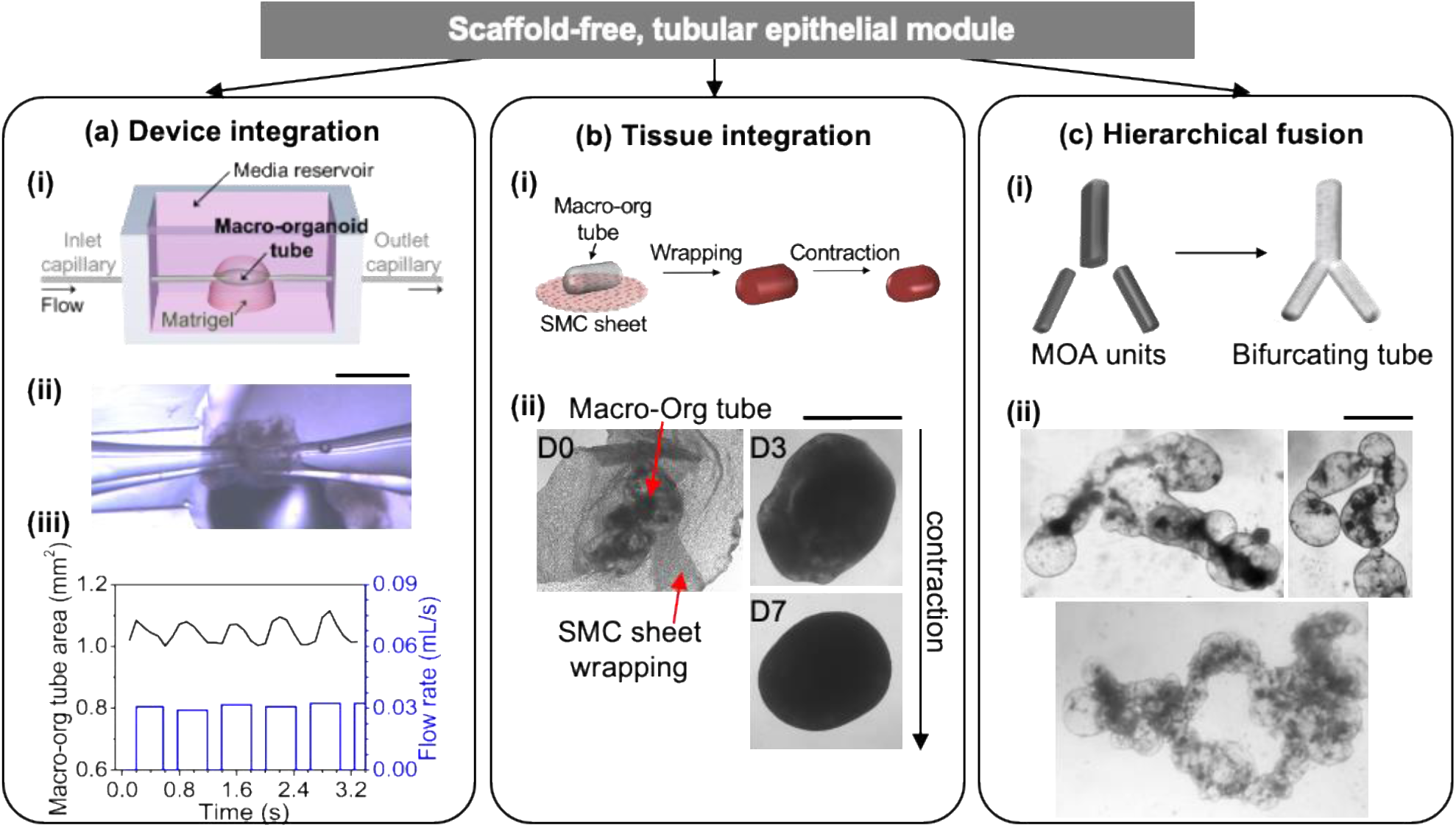
Engineered MOA tubes as scaffold-free building modules. a. Application in device-integration for a flow-able MOA tube. (i) Two pulled glass pipettes cannulate the MOA tube and connect its lumen to an external perfusion pump. (ii) Representative image (4 independent experiments) of a MOA tube under media perfusion. Scale bar, 1mm. (iii) Quantification of the MOA lumen area (black curve) in response to a peristaltic input flow (blue curve). b, Application in tissue integration. (i) Schematic representation of the SMC-MOA tube co-culture procedure to form an integral tissue construct mimicking the mouse trachea. (ii) Representative images (3 independent experiments) of the co-culture. Scale bar, 1mm. c, Biofabrication via hierarchical MOA fusion. (i) Schematic representation of the assembly and fusion of MOA building blocks, to reconstitute typical hieratical architecture of branched tubular organs. (ii) Representative images (3 independent experiments) of engineered bifurcating epithelial macro-tubes. Scale bars, 1mm.

## 3. Discussion and Conclusion

Classical bio-assembly processes largely rely on solid hydrogel matrices^[30],[31],[32]^ or patterned scaffolds^[5]^ to both define and maintain the architecture of engineered tissues. Recent work demonstrated that a suspension culture, in place of solid matrices, improves the homogeneity and throughput of individual organoid culture^[33]^; however, the sequential self-organizing cellular events remained unexplored. Here we expand the potential of the floating culture within an organoid-assembly workflow, by developing the MOrPF process that fused individual, heterogeneous organoids into tissue-scale epithelial tubes of defined geometry. Such efficient organoid fusion is realized in a Matrigel-depleted, free-floating system that harnesses the self-organization capacity of organoids. Importantly, we discovered two critical fusion steps in MOA fusion and lumenization: inter-organoid surface integration and subsequent inner matter release. Interestingly, our findings showed great morphological similarities to selected *in vivo* epithelial tubulogenesis^[34]^. The establishment of inter-organoid envelopes in MOrPF mirrors the formation of protrusion-like fusion fronts between adjacent tubular branches in the embryonic *Drosophila* trachea^[35]^ and chicken lung^[34]^. The release of inner cell matter from MOAs may resemble cavitation (elimination of redundant cells from a solid-core tissue) in the developing *Drosophila* wing^[36]^ and mouse salivary gland^[37]^. It would be of interest to further investigate the mechanisms of inter-organoid adhesion and cell migration^[38]^ during envelopment, and the possible involvement of apoptosis^[36],[39]^ during inner cell deposition and release.

Moreover, our study suggests an alternative design pathway for epithelial organoid engineering, in which shape-patterning via molding is required only at the initial organoid assembly stage, while the prescribed geometry can be retained long-term in the floating culture. This is in stark contrast to existing strategies of constructing tubular epithelial structures, which rely on exogenous shape-supporting matrices or scaffolds^[17],[40]^. Theoretical modelling suggests that the long-term shape stability of patterned MOAs originates from the biomechanical properties of fluid-filled epithelial organoid cysts. This also makes the fusion mechanism of cystic organoids distinct from that of solid-core spheroids, in which shape-maintenance is either transient (days), or requires a bounding scaffold or matrix^1,4,5^. Our engineered MOA tubes not only reproduce the elongated geometry and hollow characteristic of epithelial organs, but are upscale-able to match the dimension of the mouse tissues. Interestingly, they reach such macroscopic scale by employing a lumenization strategy during organoid self-organization, while circumventing the diffusion limit and vasculature requirement faced by the thick tissue manufacturing via the fusion of solid core aggregates. In summary, MOrPF engineered epithelial tubes are controllable and reproducible in geometry, readily accessible for further co-culture or matrix integration, and compatible with fluidic techniques to enable lumen access and fluid transport. We envisage that such macroscopic tubular epithelial building blocks could have broad implications in creating size-relevant, structurally complex, multitissue organ mimics^[14]^.

## 4. Experimental section

### Generation and culture of mouse airway organoids

Mouse airway organoids were generated following a published protocol^[41]^. Experiments were approved by local ethical review committees and conducted according to UK Home Office project license PC7F8AE82. Mice were bred and maintained under specific-pathogen-free conditions at the Cambridge Stem Cell Institute and Gurdon Institute of University of Cambridge. Briefly, trachea was dissected from 7-13 weeks old mice, and incubated in 50 U/mL dispase (Sigma) for 45 minutes (37°C). 10 mL PBS was flushed through the trachea using a syringe needle to release the trachea epithelium. Extracted cell sheets were incubated in trypsin for 5 minutes (37 °C) and dissociated single cells were mixed with growth factor-reduced Matrigel (Corning) on ice to a concentration of 1 x 10^5^ cells/mL. 30 μL dropets of Matrigel-cell mixture was plated in a 48-well plate and left in incubator for 15 minutes (37°C) for gelation. Matrigel droplets were topped with 250 μL DMEM-Ham’s F-12 media supplemented with 0.5 mg/mL penicillin-streptomycin, 10 μg/mL insulin, 5 μg/mL transferrin, 0.1 μg/mL cholera toxin, 25 ng/mL EGF, 30 μg/mL bovine pituitary extract, 5% FBS and 0.01 μM retinoic acid. Media was exchanged every 2 days.

### Design and fabrication of MOA shape-pattering molds

Negative Polylactic acid (PLA) molds with convex, tubular features (1-5 mm in length, 0.8-1.5 mm in width, 1 mm in height) were designed in Autodesk Inventor Professional 2018 software and printed with an Ultimaker 3D printer. PDMS (Sylgard 184) and curing agent is mixed at 10:1 ratio and degassed in a desiccator for 1 hour. The mixture was poured onto the PLA mold and placed in an oven at 60 °C for 6 hours. After curation, PDMS containing arrays of concave wells was peeled from PLA and immersed in ethanol for 18 hours to remove uncured PDMS polymers. The PDMS mold was then autoclaved and immersed in an anti-cell adherence solution (STEMCELL, Catalogue 07010) for 1 hour. After removal of the anti-adherence coating, PDMS molds were washed with PBS three times and to dry for use as a template for MOA shape pattering.

### MOA patterning via molding

Day-12 organoids were harvested from Matrigel and centrifuged at 500 x *g* for 5 minutes (0°C). Supernatant was discarded and organoid pellet was resuspended with cold media (0°C), to remove residual Matrigel on the organoid surface. Organoids underwent a second centrifugation at 500 x *g* for 5 minutes (0°C) to form a compact pellet. Supernatant was removed and organoid pellet was carefully pipetted into the PDMS shape-patterning wells until the concave space was filled. To recreate mouse trachea anatomy (1-1.5 mm inner diameter), airway organoids were assembled within tubular wells of the same dimension (1-1.5 mm in width, 2-6 mm in length, 1 mm in height). MOAs were then overlaid with media and incubated in PDMS molds overnight to acquire the prescribed shape via contact guidance. Subsequently, MOAs were released from PDMS molds and cultured in a media-only condition in ultra-low attachment 6-well plates (Corning).

### MOA formation via pipette extrusion

Day-12 organoids were harvested from Matrigel and centrifuged at 500 x *g* for 5 minutes (0°C). Supernatant was discarded and organoid pellet was resuspended with cold media (0°C), to remove residual Matrigel on the organoid surface. Organoids underwent a second centrifugation at 500 x *g* for 5 minutes (0°C) to form a compact pellet. Upon supernatant removal, organoid pellet was sucked into a 200 μL pipette tip and manually extruded into a string-like shape in warm media. The string-shaped aggregates were cultured long-term in a media-only, ultra-low attachment 6-well plate (Corning).

### Image acquisition and analysis

Bright-field images of MOAs were acquired daily using an Olympus optical microscope with 4x and 10x objectives. The projected area of MOAs was measured using ImageJ software. For AIMR (= MOA area/inner matter area) quantification, optical images were corrected with background subtraction and converted into binary masks using the threshold function in ImageJ.

Time-lapse imaging was acquired by using a confocal spinning-disk microscope system (Intelligent Imaging Innovations, Inc. 3i). The imaging system was composed of an Observer Z1 inverted microscope (Zeiss), a CSU X1 spinning disk head (Yokogawa), and a Quant EM 512SC camera (Photometrics). Silicon micro-inserts (4 well, Ibidi) were inserted into each well of an uncoated 8-well μ-slide (Ibidi). 8-12 Matrigel-depleted organoids were positioned within close proximity (less than 80 μm) in each mini-compartment created in the μ-slide. Organoids were grouped into three culture conditions: Floating, Gel-suspension (culture media supplemented with 20% volume fraction Matrigel in suspension) and the conventional Matrigel droplet culture. The μ-slide was mounted in a mini imaging compartment with controlled temperature (37 °C) and CO_2_ concentration (5%). Organoid behaviours were recorded every 2 days with a 10x objective over a 10-day observation period. A x, y, z scanning mode was used to obtain z stack images. 3D image reconstruction and quantification of organoid dynamics (inter-organoid gap, envelope length, projected area of released inner matter) were performed in Slidebook6 software.

### Immunofluorescent and histological staining

For immunofluorescent study, MOAs were prepared *via* pipette extrusion. MOAs at early-lumenization, mid-lumenization and stabilization stages were fixed with 4% paraformaldehyde (PFA) for 1 hour and washed three times with PBS under a fume hood. For immunofluorescent staining, fixed samples were permeabilized in 0.2% Triton X-100 in PBS for 1 hour, then blocked in 2% normal donkey serum (NDS) in PBS for 1 hour at room temperature. Primary antibodies (in 5% NDS/PBS solution at 1:200) were added and left overnight at 4°C, then washed off with 0.2% Tween-20 in PBS. Secondary antibodies were (in PBS at 1:2000) incubated for 1 hour at room temperature, then washed off with 0.2% Tween-20 in PBS. For staining DNA, DAPI (1:1000 dilution) was used for 5 minutes at room temperature. Antibodies are listed in the Supporting Information.

For histological staining, fixed samples were embedded in paraffin wax and sectioned on a xyz microtome. Hematoxylin and eosin were used as dyes to stain respectively the cell nucleus (blue) and cytoplasm (pink).

### Design and fabrication of a perfusion chip for MOA tubes

The millifluidics perfusion device was composed of a single-channel PDMS chip and a pair of pulled glass capillaries (1 mm outer diameter). A PLA template for the PDMS chip was designed in Autodesk inventor 2018 and printed with an Ultimaker 3D printer. A syringe needle (1 mm outer diameter) was assembled in the PLA construct, before pouring the PDMS-curing agent mixture (10:1) into the PLA mold. After curing (60 °C, 6 hours), the needle was removed and PDMS was peeled from the PLA, followed by an immersion in ethanol for 18 hours. A 4 mm hole was punched in the centre of the PDMS chip, as a space to accommodate the MOA tube. The PDMS chip was then bonded with a thin glass slide using a plasma cleaner (Harrick plasma) and autoclaved before use.

A MOA tube was placed in the central culture area and Matrigel was added to fill the remaining culture space. The chip was incubated for 15 minutes to allow Matrigel gelation, followed by a top-up of the media reservoir with warm media. The airway-on-a-chip was stabilized in the incubator for at least 6 hours before mounted on an Olympus SZX16 upright optical microscope. Two glass capillaries were prepared using a micropipette puller (Sutter Instrument) and cut into 50 μm tip diameter with a micro forge (Narishige). Capillaries were autoclaved and pre-filled with culture media before insertion through the two ports of the PDMS chip. The inlet and outlet capillaries were connected to a Fisherbrand peristaltic pump through a media pre-filled silicone tubing, and carefully advanced toward the MOA tube to cannulate the lumen. Direction and velocity of the media flow were tuned by controlling the rotating direction (clockwise or anti-clockwise) and speed (level 1 to 10) of the rotors. Flow rate calibration was performed prior to perfusion by recording the weight change of media droplets flowing from the silicone tubing.

### Engineered co-culture of SMCs and MOA tubes

Mouse vascular aortic smooth muscle cells MOVAS (ATCC^®^ CRL-2797) were purchased from ATCC and cultured in DMEM (DB) supplemented with 10% FBS and antibiotics. To form SMC sheets, SMCs were passaged on 80% confluence and incubated in a temperature-responsive Nunc™ Dishe with UpCell™ Surface (ThermoFisher). When cells reached confluence, the UpCell plate was transferred to room temperature (20°C) to initiate temperature-regulated cell sheet detachment. SMC medium was discarded, and a MOA tube was placed atop of a SMC sheet. Subsequently, the SMC sheet was careful lifted from the well bottom using forceps and wrapped around the MOA tube, to create the co-culture construct. Organoid media was then added to the assembled tissue, which was cultured for 5 days before antibody staining.

### Statistical analysis

All statistical analyses were performed in OriginLab 2015 software. *P* values were calculated using two-tailed Student’s t-test and Mann-Whitney U test. *P*-values less than 0.05 were considered statistically significant. All box plots extend from the 25th to 75th percentiles, with a line at the median and whiskers extending to maximum and minimum data points. Each experiment was repeated at least twice using independent batches of organoids.

## Supporting Information

Supporting Information is available from the Wiley Online Library or from the author.

## Acknowledgements

We thank Prof. Kim Dora, from Department of Pharmacology, University of Oxford for advice on vessel perfusion; Mr Lukas Vasadi, from Department of Engineering, University of Cambridge, for technical supports on the perfusion set-up. This work was supported by the European Research Council (ERC-StG, 758865; ERC-StG, 679411) and the Wellcome and the Royal Society (107633/Z/15/Z). YL. was a recipient of Schlumberger Faculty for the Future scholarship and Trinity Henry-Barlow scholarship. C.D. was supported by the Cancer Research UK Cambridge Centre PhD Studentship. Y.L., J-H.L. and Y.Y.S.H. conceived and designed experiments.

## References

[1] K. Jakab, A. Neagu, V. Mironov, R. R. Markwald, G. Forgacs, Proc. Natl. Acad. Sci. U. S. A. 2004, 101, 2864.

[2] S. V. Murphy, A. Atala, Nat Biotech 2014, 32, 773.

[3] M. A. Skylar-Scott, S. G. M. Uzel, L. L. Nam, J. H. Ahrens, R. L. Truby, S. Damaraju, J. A. Lewis, Sci. Adv. 2019, 5, eaaw2459.

[4] V. Mironov, R. P. Visconti, V. Kasyanov, G. Forgacs, C. J. Drake, R. R. Markwald, Biomaterials 2009, 30, 2164.

[5] A. P. Rago, D. M. Dean, J. R. Morgan, Biotechnol. Bioeng. 2009, 102, 1231.

[6] A.-C. Tsai, Y. Liu, X. Yuan, T. Ma, Tissue Eng. Part A 2015, 21, 1705.

[7] D. G. Belair, C. J. Wolf, C. Wood, H. Ren, R. Grindstaff, W. Padgett, A. Swank, D. MacMillan, A. Fisher, W. Winnik, B. D. Abbott, PLoS One 2017, 12, e0184155.

[8] Y. Takeoka, K. Matsumoto, D. Taniguchi, T. Tsuchiya, R. Machino, M. Moriyama, S. Oyama, T. Tetsuo, Y. Taura, K. Takagi, T. Yoshida, A. Elgalad, N. Matsuo, M. Kunizaki, S. Tobinaga, T. Nonaka, S. Hidaka, N. Yamasaki, K. Nakayama, T. Nagayasu, PLoS One 2019, 14, e0211339.

[9] B. Ayan, D. N. Heo, Z. Zhang, M. Dey, A. Povilianskas, C. Drapaca, I. T. Ozbolat, Sci. Adv. 2020, 6, eaaw5111.

[10] T. Sato, R. G. Vries, H. J. Snippert, M. van de Wetering, N. Barker, D. E. Stange, J. H. van Es, A. Abo, P. Kujala, P. J. Peters, H. Clevers, Nature 2009, 459, 262.

[11] K. W. McCracken, E. M. Catá, C. M. Crawford, K. L. Sinagoga, M. Schumacher, B. E. Rockich, Y.-H. Tsai, C. N. Mayhew, J. R. Spence, Y. Zavros, J. M. Wells, Nature 2014, 516, 400.

[12] M. Fujii, T. Sato, Nat. Mater. 2020, DOI 10.1038/s41563-020-0754-0.

[13] J. A. Brassard, M. P. Lutolf, Cell Stem Cell 2019, 24, 860.

[14] T. Takebe, J. M. Wells, Science (80-.). 2019, 364, 956 LP.

[15] Y. Xiang, Y. Tanaka, B. Patterson, Y.-J. Kang, G. Govindaiah, N. Roselaar, B. Cakir, K.-Y. Kim, A. P. Lombroso, S.-M. Hwang, M. Zhong, E. G. Stanley, A. G. Elefanty, J. R. Naegele, S.-H. Lee, S. M. Weissman, I.-H. Park, Cell Stem Cell 2017, 21, 383.

[16] J. A. Bagley, D. Reumann, S. Bian, J. Lévi-Strauss, J. A. Knoblich, Nat. Methods 2017, 14, 743.

[17] N. Sachs, Y. Tsukamoto, P. Kujala, P. J. Peters, H. Clevers, Development 2017, 144, 1107 LP.

[18] J. R. Rock, M. W. Onaitis, E. L. Rawlins, Y. Lu, C. P. Clark, Y. Xue, S. H. Randell, B. L. M. Hogan, Proc. Natl. Acad. Sci. 2009, 106, 12771 LP.

[19] T. A. Sebrell, B. Sidar, R. Bruns, R. A. Wilkinson, B. Wiedenheft, P. J. Taylor, B. A. Perrino, L. C. Samuelson, J. N. Wilking, D. Bimczok, Cell Tissue Res. 2018, 371, 293.

[20] C. J. Chan, M. Costanzo, T. Ruiz-Herrero, G. Mönke, R. J. Petrie, M. Bergert, A. Diz-Muñoz, L. Mahadevan, T. Hiiragi, Nature 2019, 571, 112.

[21] T. P. J. Wyatt, J. Fouchard, A. Lisica, N. Khalilgharibi, B. Baum, P. Recho, A. J. Kabla, G. T. Charras, Nat. Mater. 2020, 19, 109.

[22] M. Doi, Soft Matter Physics, Oxford University Press, 2013.

[23] R. Simson, E. Wallraff, J. Faix, J. Niewöhner, G. Gerisch, E. Sackmann, Biophys. J. 1998, 74, 514.

[24] E. Sackmann, R. F. Bruinsma, ChemPhysChem 2002, 3, 262.

[25] J. Y. Co, M. Margalef-Català, X. Li, A. T. Mah, C. J. Kuo, D. M. Monack, M. R. Amieva, Cell Rep. 2019, 26, 2509.

[26] D. Huh, B. D. Matthews, A. Mammoto, M. Montoya-Zavala, H. Y. Hsin, D. E. Ingber, Science (80-.). 2010, 328, 1662 LP.

[27] I. Orhon, N. Dupont, M. Zaidan, V. Boitez, M. Burtin, A. Schmitt, T. Capiod, A. Viau, I. Beau, E. W. Kuehn, G. Friedlander, F. Terzi, P. Codogno, Nat. Cell Biol. 2016, 18, 657.

[28] M. Navarro, J. Ruberte, A. Carretero, in (Eds.: J. Ruberte, A. Carretero, M.B.T.-M.M.P. Navarro), Academic Press, 2017, pp. 147–178.

[29] S. E. Park, A. Georgescu, D. Huh, Science (80-.). 2019, 364, 960 LP.

[30] O. C. Tysoe, A. W. Justin, T. Brevini, S. E. Chen, K. T. Mahbubani, A. K. Frank, H. Zedira, E. Melum, K. Saeb-Parsy, A. E. Markaki, L. Vallier, F. Sampaziotis, Nat. Protoc. 2019, 14, 1884.

[31] M. M. Capeling, M. Czerwinski, S. Huang, Y.-H. Tsai, A. Wu, M. S. Nagy, B. Juliar, N. Sundaram, Y. Song, W. M. Han, S. Takayama, E. Alsberg, A. J. Garcia, M. Helmrath, A. J. Putnam, J. R. Spence, Stem Cell Reports 2019, 12, 381.

[32] N. Gjorevski, N. Sachs, A. Manfrin, S. Giger, M. E. Bragina, P. Ordóñez-Morán, H. Clevers, M. P. Lutolf, Nature 2016, 539, 560.

[33] N. Brandenberg, S. Hoehnel, F. Kuttler, K. Homicsko, C. Ceroni, T. Ringel, N. Gjorevski, G. Schwank, G. Coukos, G. Turcatti, M. P. Lutolf, Nat. Biomed. Eng. 2020, DOI 10.1038/s41551-020-0565-2.

[34] M. A. Palmer, C. M. Nelson, Dev. Dyn. 2020, n/a, DOI 10.1002/dvdy.215.

[35] L. Gervais, G. Lebreton, J. Casanova, Dev. Biol. 2012, 362, 187.

[36] E. Moreno, K. Basler, G. Morata, Nature 2002, 416, 755.

[37] A. S. Tucker, Semin. Cell Dev. Biol. 2007, 18, 237.

[38] D. Sutherland, C. Samakovlis, M. A. Krasnow, Cell 1996, 87, 1091.

[39] J. Debnath, K. R. Mills, N. L. Collins, M. J. Reginato, S. K. Muthuswamy, J. S. Brugge, Cell 2002, 111, 29.

[40] J. A. Reid, P. A. Mollica, R. D. Bruno, P. C. Sachs, Breast Cancer Res. 2018, 20, 122.

[41] J.-H. Lee, T. Tammela, M. Hofree, J. Choi, N. D. Marjanovic, S. Han, D. Canner, K. Wu, M. Paschini, D. H. Bhang, T. Jacks, A. Regev, C. F. Kim, Cell 2017, 170, 1149.

