## Supplementary Information for "Bioassemblying Macro-Scale, Lumnized Airway Tubes of Defined Shape via Multi-Organoid Patterning and Fusion"

**Supplementary Figures**

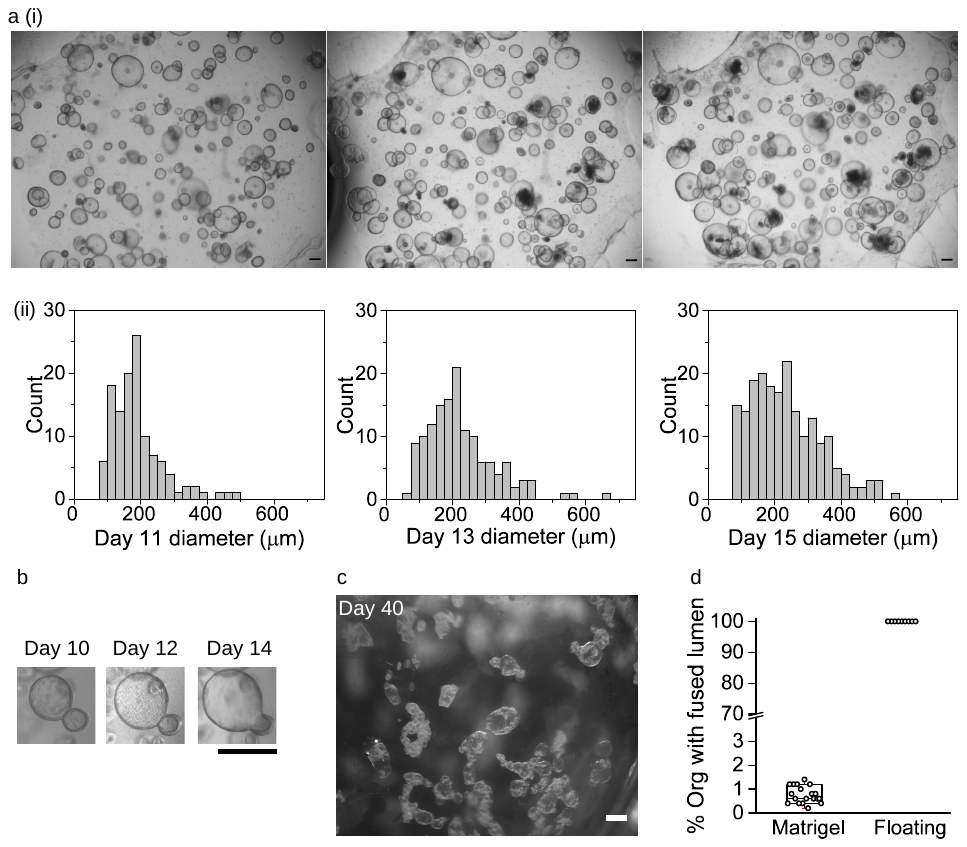

**Figure S1.** Mouse airway organoids grow to heterogeneous sizes and occasionally fuse in Matrigel. a, Representative images (3 independent experiments) of the developing organoids (i) and their size distribution (ii) over time in Matrigel. Scale bars, 200 μm. b, Representative images of occasional organoid fusion in Matrigel. Scale bar, 500 μm. c, Representative images (3 independent experiments) of spontaneous organoid aggregation and fusion in the Floating culture, when no preliminary shape-patterning was applied. Scale bar, 1 mm. d, Percentage of fused organoids in Matrigel and the floating culture. Matrigel droplets, n=19. Floating, n=20.

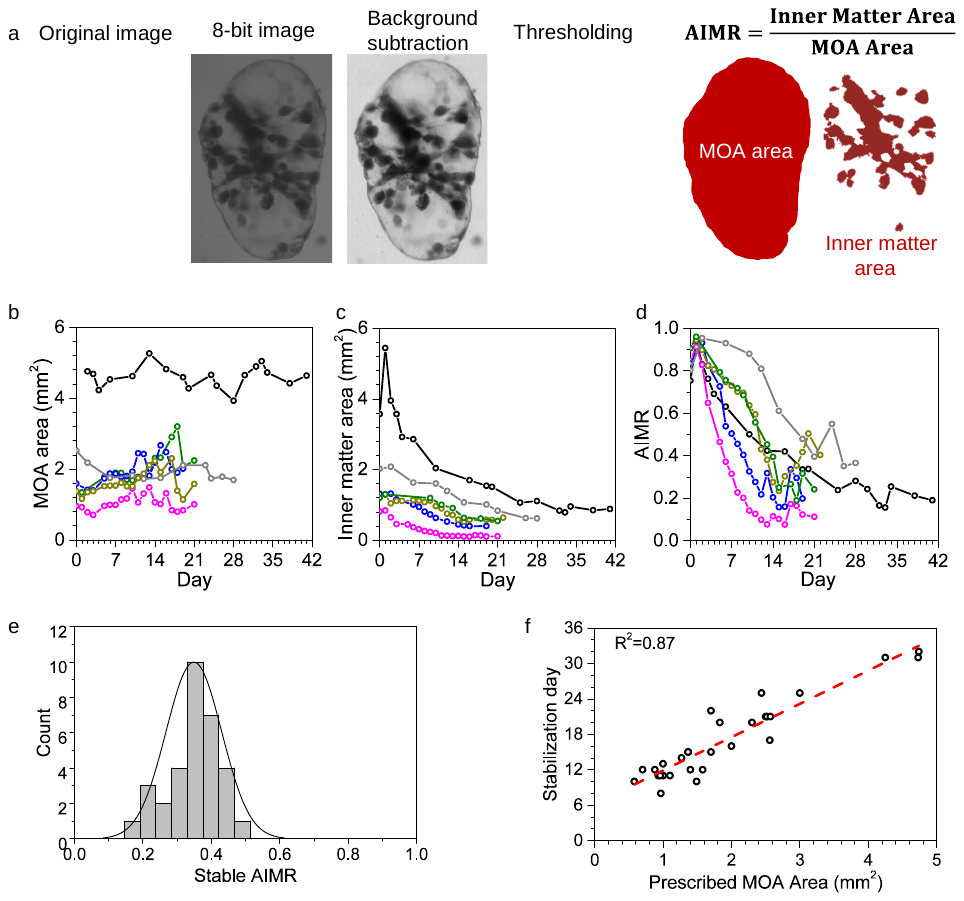

**Figure S2**. Macroscopic characterization of MOA morphology during MOrPF. a, Schematic representation of AIMR definition and quantification. Bright field image of the MOA undergoes three sequential image processing steps, to obtain a binary mask for AIMR quantification. b-d, Dynamics of MOA area, inner matter area and AIMR for MOAs of different sizes. Each curve represents one MOA. n=6. e, Quantification of the stable AIMR value for fused MOAs with various sizes. n=29. Black curve represents the normal distribution fit. f, Days required to achieve MOA stabilization as a function of MOA size. n=29.

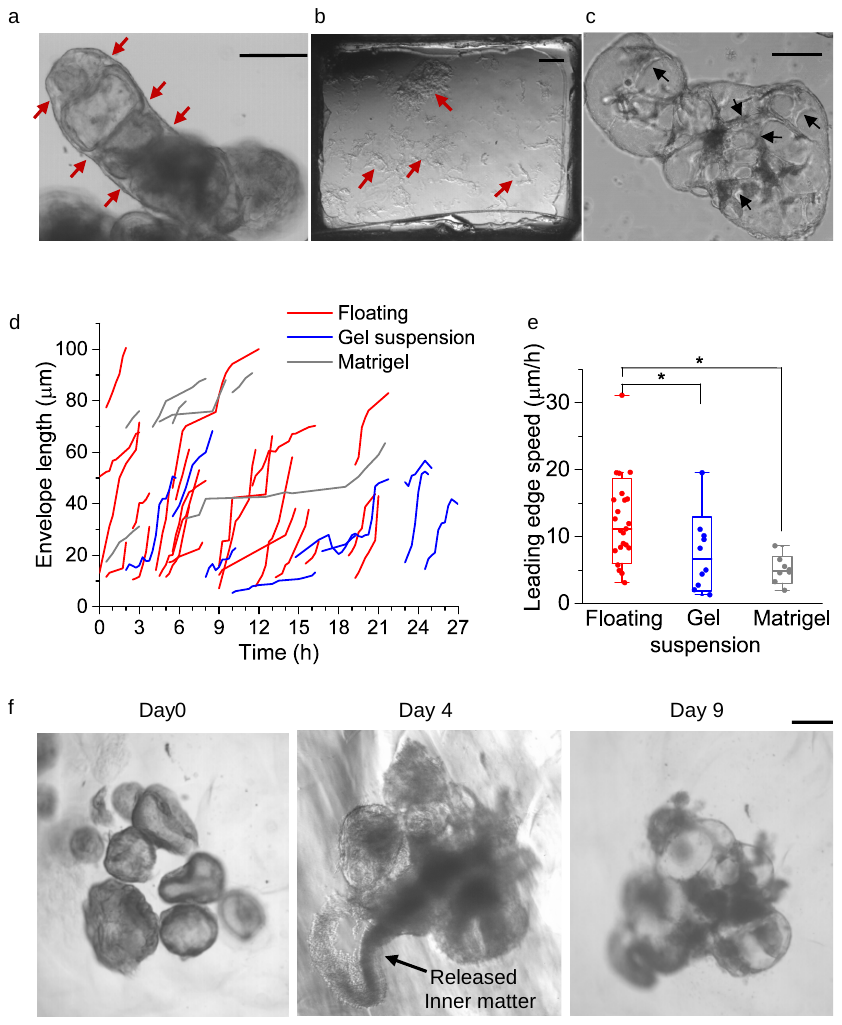

**Figure S3.** Microscopic characterization of MOA fusion dynamics. a, Representative image (3 independent experiments) of inter-organoid envelopes developed at multiple locations (red arrows) on a MOA. Scale bar, 200 μm. b, Representative image (3 independent experiments) of Gel suspension containing suspending Matrigel fragments (red arrows). Scale bar, 200 μm. c, Representative image (3 independent experiments) of a developing MOA with ‘interfacial disks’ (black arrows). Scale bar, 200 μm. d,e, Quantification of the length (d) and leading edge speed (e) of inter-organoid envelopes in different culture conditions. Each curve represents one developing envelope. Midline = median, box = 25th-75th percentiles, Whisker = min and max values. Floating, n = 23. Gel suspension, n = 8. Matrigel, n = 23. *P <0.005 by Student’s t-test. f, Representative images (3 independent experiments) of MOA fusion with hindered inner matter release in agarose suspension (0.5% weight percent in media). Scale bar, 250 µm.

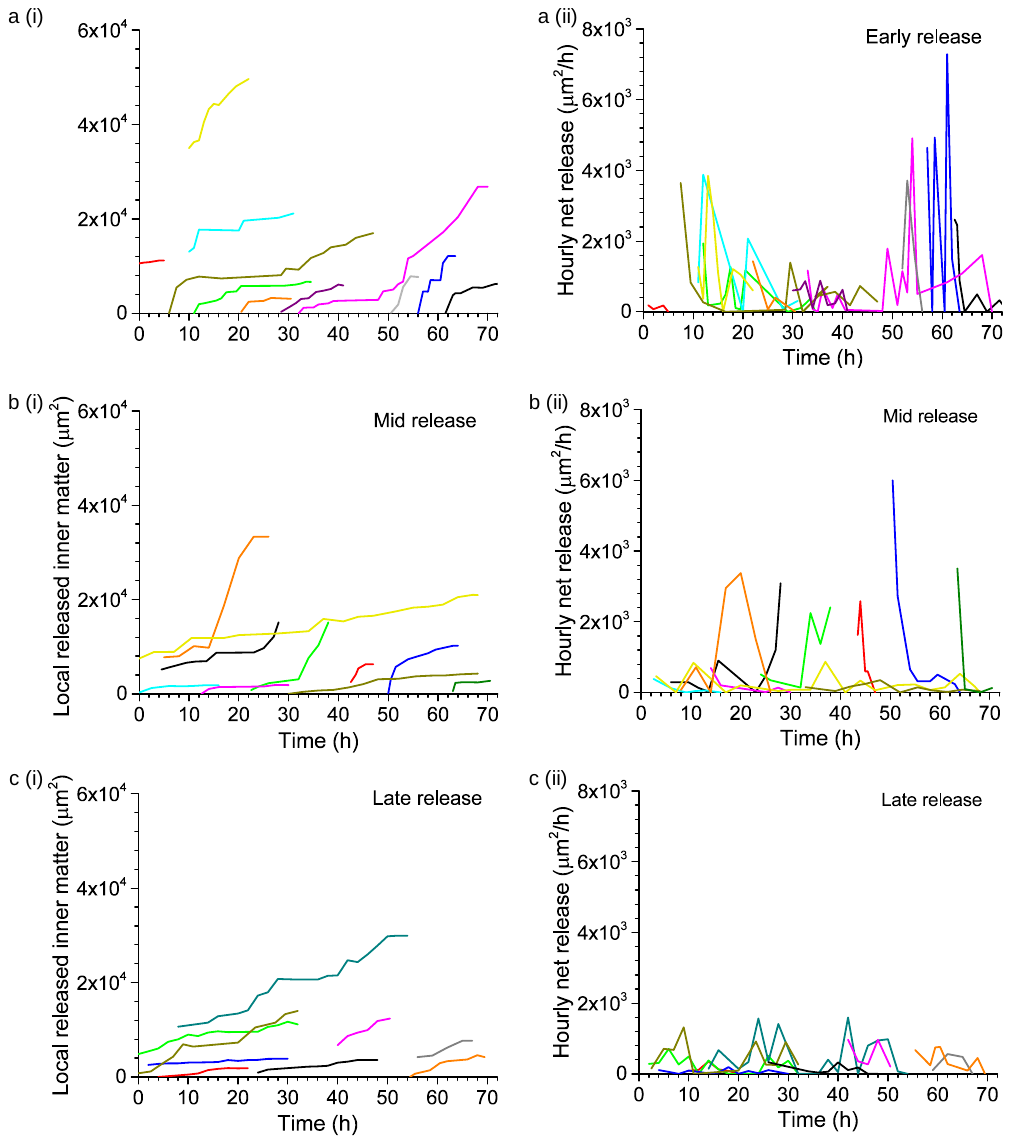

**Figure S4.** Dynamics of MOA inner matter release in the Early (a), Mid (b) and Late (c) lumenization phases. (i) Area dynamics of released inner matter. Each curve represents one local release site on a MOA. Early, n=11; Mid, n=10; Late, n=9. (ii) Calculated hourly release speeds for corresponding release sites shown in the left panel.

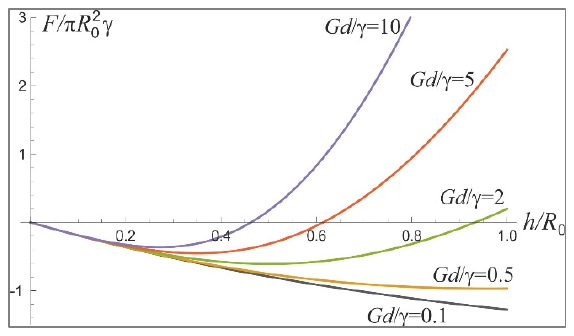

**Figure S5.** Theoretical model predicts the normalized energy potential as a function of normalized organoid deformation(*h/R_0_*) for different *Gd/γ*.

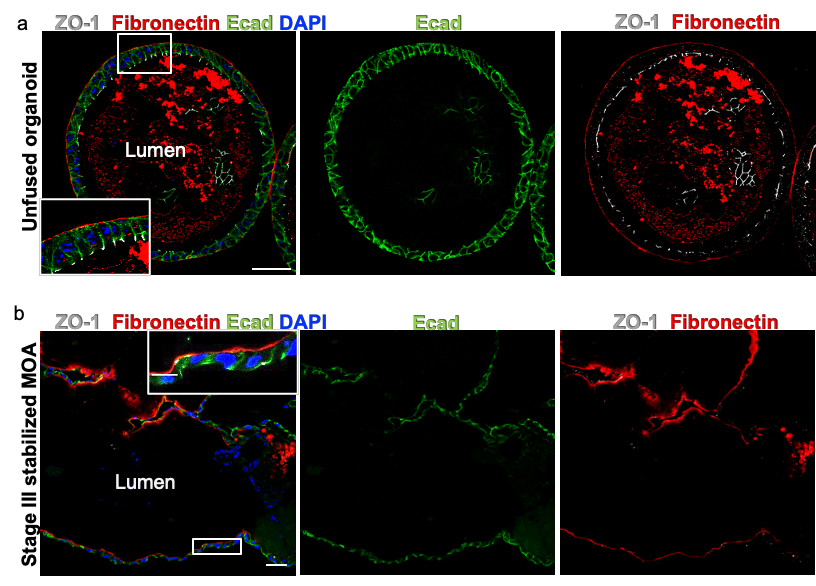

**Figure S6.** Representative immunostaining (2 independent experiments) for polarity markers in unfused control airway organoids (a) and stabilized MOAs-Stage III (b). a, Wholemount IF staining of a control airway organoid showing basal localization of fibronectin (red), apical localization of tight-junction protein ZO-1 (white) and lateral localization of epithelial Ecad (green). Scale bar, 30 μm. b, Paraffin sections of a Stage III stabilized MOA showing apical localization of fibronectin (red) suggestive of polarity inversion, maintenance of Ecad expression and downregulation of ZO-1. DAPI (blue) stains nuclei. Insets show magnified version of the boxed regions. Scale bar, 30 μm.

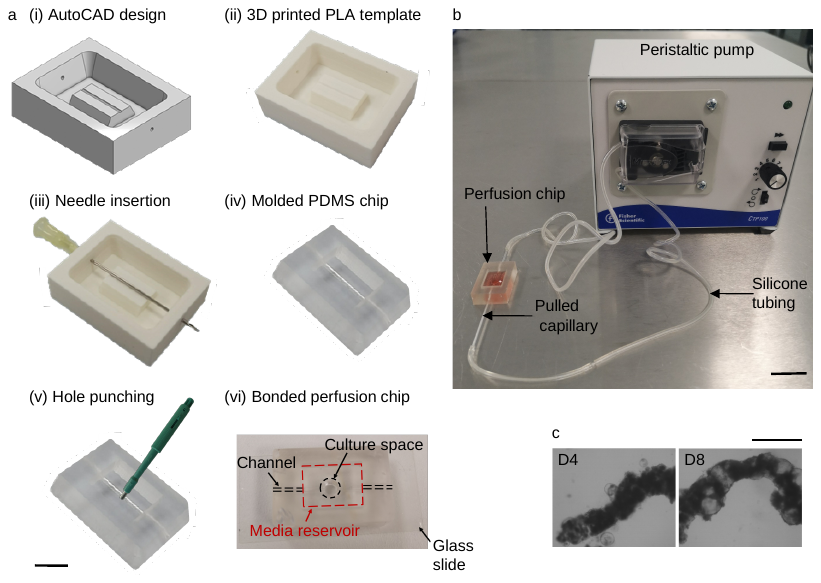

**Figure S7**. Perfusion setup (a, b) for the flow-able MOA tubes and the biofabrication of a pseudo mouse intestinal tube (c). a, Fabrication procedure of a PDMS perfusion chip. Scale bar, 10 mm. b, Flow system for the MOA tubes. Scale bar, 3 cm. c, Representative image (2 independent experiments) of a mouse intestinal MOA tube created via manual extrusion in the floating condition. Scale bar. 1mm.

**Theoretical modelling**

It is assumed that at the organoid’s reference state with diameter *R_0_*, the elastic shells are approximately tension-free, as any initial tension in this epithelial layer caused by Laplace pressure under the curved surface had enough time to relax via the plastic flow of epithelial cells (which takes about 1 day, based on the migration speed ~10 µm/hr indicated in Figure S3e, Supporting information, over an organoid diameter of 200 µm). After equilibration, the shape of such a single organoid is maintained by the high bending rigidity of the epithelial layer: given the elastic modulus of the layer as *G* = 200 Pa^21^ and its thickness *d* = 20 mm (Figure 5a), the bending modulus (often called the flexural rigidity of a membrane) is equal to $B = \frac{1}{9}Gd^{3} = {10}^{8} \text{kJ/mol}$^42^, which is much higher than the characteristic energy of thermal fluctuations ($k_{B}T=2.4 \text{kJ/mol}$), making the curved organoid wall effectively stiff.

In the formation of the fused organoid shape, two physical effect are competing: there is an energy gain by increasing the area of the ‘adhesion disk’ due to the favorable surface energy (often called surface tension in the studies of wetting), and an elastic energy penalty in the stretched organoid shells. The estimate of the surface tension, *γ* = 2 mN/m, reflecting the cell-cell adhesion strength in two shells in contact, is reasonable considering it is close to tension for epithelial cadherin-mediated adhesion (estimated to be 3 mN/m^43,44^). Note that strong cell-cell adhesion could be established in the first 10 minutes after the initial contact^45^, before the full adhesion complex is assembled and matured. In contrast, the lower limit of surface tension is due to the non-specific cell-substrate adhesion (0.01 mN/m), which is only relevant in the first few minutes after contact^46^.

Since each original organoid is deformed into a spherical cap, at constant internal volume on the time scale for stable establishment of the dumbbell shape (~hours), its bilayer shell becomes stretched and this costs elastic energy. Thus, the spherical caps in Figure 3c(i) are of a slightly inflated radius *R*>*R_0_*. The inflating ratio *λ*= *R*/*R_0_* is therefore a measure of the elastic deformation in the membrane, causing the elastic penalty well studied in the theory of rubber balloons^22^. Note that the inflating ratio is independently related to the shape of the spherical caps in the deformed spheres, since we assume that at such short time scale, the inner volume must remain constant. As a result of this volume conservation constraint, the relationship reads:

$$\frac{h}{R_{0}}\approx\lambda-1+\frac{2\sqrt{\lambda^{3}-1}}{\sqrt{3}}$$

The combined energy is expressed as a function of the overlapped height *h* and the inflating ratio *λ*, which is related to *h* by the constraint above.

$$F\left( h,\lambda\right)=2\pi R_{0}^{2}\left( 2-\frac{h \lambda}{R_{0}}\left( 1-\frac{h}{2\lambda R_{0}} \right) \right)d\cdot G\left( 2\lambda^{2}+\frac{1}{\lambda^{4}}-3 \right)-2\pi R_{0}h \lambda\left( 1-\frac{h}{2\lambda R_{0}} \right)\cdot\gamma$$

In the first term, we have the elastic energy penalty in the stretched shell of each organoid (a membrane of elastic modulus *G*), with the area of the outer spherical cap: $2\pi R_{0}^{2}\left( 2-\frac{h \lambda}{R_{0}}\left( 1-\frac{h}{2\lambda R_{0}} \right) \right)$. The second term is the adhesion energy gain over the adhesion disk area: $2\pi R_{0}h \lambda\left( 1-\frac{h}{2\lambda R_{0}} \right)$. Importantly, the resulting shape of the fused organoid is determined by a single non-dimensional factor, which is the ratio of the elastic and adhesion parameters: $Gd/\gamma$.

**List of antibodies**

Primary antibodies:

| **Antibody** | **Clone** | **Manufacturer** | **Catalogue Number** |
| --- | --- | --- | --- |
| Keratin-5 (Rabbit) | Poly19055 | Biolegend | 905501 |
| Acetyl-tubulin (Mouse) | 6-11B-1 | Sigma | T7451 |
| E-cadherin (Rat) | ECCD-2 | Life Tech | 131900 |
| Krt8 (Rat) | TROMA-I | DSHB | TROMA-I |
| ZO1 (Rabbit) | Polyclonal | Invitrogen | 40-2200 |
| Fibronectin (Sheep) | Polyclonal | R&D | AF1918 |
| p63-α (D2K8X) Rabbit | Monoclonal | Cell Signaling | 13109 |

Secondary antibodies:

| **Antibody** | **Class** | **Manufacturer** | **Catalogue Number** |
| --- | --- | --- | --- |
| Donkey anti-Mouse IgG (H+L)  Alexa Fluor 488 | Polyclonal | Thermofisher Scientific | A-21202 |
| Donkey anti-Rabbit IgG (H+L)  Alexa Fluor 488 | Polyclonal | Thermofisher Scientific | A-21206 |
| Donkey anti-Rabbit IgG (H+L)  Alexa Fluor 555 | Polyclonal | Thermofisher Scientific | A-31572 |
| Donkey anti-Rat IgG (H+L)  Alexa Fluor 488 | Polyclonal | Thermofisher Scientific | A-21208 |
| Goat anti-Rat IgG (H+L)  Alexa Fluor 647 | Polyclonal | Thermofisher Scientific | A-21247 |

**Video S1.** Inter-organoid gap closure and envelope development. This video shows the movement of one organoid towards an adjacent organoid, for closely-spaced organoids cultured in the floating environment. Once in close contact, the organoid sends out a cell envelope towards its neighboring organoid. Inter-organoid envelopes keep extending on the surface of the adjacent organoid and can emerge at multiple locations, until all organoids in proximity integrate into one MOA.

**Video S2.** MOA compaction. This video shows the decreasing inter-organoid space, resulting in densified organoid packaging within a MOA.

**Video S3.** Inner matter release 1. This video shows the release of inner cell materials from two local release sites on a big MOA.

**Video S4.** Inner matter release 2. This video shows the release of inner cell materials from two local release sites on a small MOA. Nucleus are labelled red.

**Video S5.** Perfusion of an airway MOA tube. This video (real-time) shows the introduction of a peristaltic media flow into the MOA lumen. Connected to an external pump by a pair of glass pipettes, the MOA can expand and shrink in response to the peristaltic perfusion.
